# Alpha activity in the ventral and dorsal visual stream controls information flow during working memory

**DOI:** 10.1101/180166

**Authors:** Marcin Leszczynski, Juergen Fell, Ole Jensen, Nikolai Axmacher

**Affiliations:** Department of Neurological Surgery, Columbia University College of Physicians and Surgeons, New York, New York, USA; Translational Neuroscience Division, Nathan Kline Institute, Orangeburg, New York, USA; Department of Epileptology, University of Bonn, Bonn, Germany; Centre for Human Brain Health, University of Birmingham, Birmingham, UK; Department of Neuropsychology, Institute of Cognitive Neuroscience, Faculty of Psychology, Ruhr University Bochum, Bochum, Germany

**Keywords:** alpha oscillations, ventral visual stream, dorsal visual stream, cross-frequency coupling, phase-amplitude coupling, working memory, face processing, cognition

## Abstract

The electrophysiological mechanisms underlying working memory maintenance of information in the ventral and dorsal visual stream (VVS, DVS) remain elusive. Here we used electrocorticography recordings covering VVS, DVS and prefrontal cortex (PFC) in epilepsy patients while they were performing a delayed match-to-sample task. The experimental conditions (face identity, orientation) were designed to engage either the VVS or DVS. Alpha power was reduced in the VVS during maintenance of face identity and in the DVS during maintenance of spatial orientation of the very same stimuli. The phase of alpha oscillations modulated broadband high-frequency activity (BHA) in both regions. Interestingly, BHA occurred across broader alpha phase ranges when task-relevant information was maintained, putatively reflecting longer excitable “duty cycles”. Our findings support a model in which the VVS and DVS are recruited by the PFC via selective reduction of alpha power. As a result, excitable duty cycles in the relevant area are extended.

## Introduction

Successful performance in a working memory (WM) task depends critically on the dynamic engagement of networks involved in representing relevant information and disengagement of networks representing any potentially distracting information. Several cortical networks important for representing information in the absence of sensory input have already been identified (Eriksson et al., 2015). Evidence from human neuroimaging has shown that the ventral (Courtney et al., 1997; Druzgal and D’Esposito, 2003; Postle et al., 2003; Ranganath et al., 2004a,b) and dorsal (Awh et al., 1999; Postle and D’Esposito, 1999; Postle et al., 2004; Postle, 2006) visual stream are two key regions involved in perception and WM maintenance of distinct visual features (e.g. face identity and spatial orientation, respectively). Studies have also identified the prefrontal cortex as critical to WM maintenance, likely providing top-down signals to the sensory cortical areas (Funahashi et al., 1989; Goldman-Rakic, 1995; Curtis and D’Esposito, 2003; Sreenivasan et al., 2014). However, the electrophysiological mechanisms underlying the contribution of ventral and dorsal visual stream to WM remain poorly understood. Similarly, the electrophysiological mechanisms supporting the communication between the prefrontal cortex and ventral/dorsal visual stream remain unclear. Cortical alpha oscillations (8-14Hz) have been associated with modulations of neural activity (Haegens et al., 2011, Spaak et al., 2012) and have been implicated in an active functional inhibition of task-irrelevant information (Jokisch and Jensen, 2007; Klimesch et al., 2007; Klimesch, 2012; Bonnefond and Jensen, 2012; Jensen et al., 2012; Payne and Sekuler, 2014; Anderson and Hanslmayr, 2015). The functional inhibition model posits that both phase and power of alpha oscillations play a role in organizing the neuronal code. Specifically, it has been suggested that the alpha phase serves to restrict active neural representations to specific phase ranges of alpha oscillations, so-called “duty cycles” (Jensen et al., 2014). Furthermore, Roux and Uhlhaas (2014) suggested that alpha-gamma phase-amplitude coupling might serve as a code for the maintenance of various sensory features (e.g. spatial information; see also Park et al., 2016) as opposed to the theta-gamma coupling suggested to support multi-item WM (Lisman and Idiart, 1995; Canolty et al., 2006; Axmacher et al., 2010b; Roux and Uhlhaas, 2014; Leszczynski et al., 2015; Daume et al., 2017). The power of alpha oscillations, in turn, may be relevant because it scales inversely with the duration of “duty cycles”, both as approximated on the level of action potentials (Haegens et al., 2011) and of high frequency activity (Osipova et al,. 2008; Spaak et al., 2012; Bonnefond and Jensen, 2015).

These effects of alpha phase and power related to functional inhibition have been demonstrated in primary sensory cortices. However, it remains unknown if phase and power of alpha activity have a similar role for functional inhibition in higher visual areas. This question is particularly important because previous research suggested that the mechanisms underlying the generation of alpha activity in the ventral visual stream differ from those in early sensory cortices (Bollimunta et al., 2008; Mo et al., 2011). In particular, alpha generators have been found in the supra-, infra- and granular layer of early visual areas (e.g. V1, V2, V4; Bollimunta et al., 2008; Haegens et al., 2015), but only in the supra- and infragranular layer of the ventral visual stream (Bollimunta et al., 2008). Whether alpha oscillations reflect functional inhibition in the ventral stream remains unknown.

Increased interareal phase coherence in the theta (Liebe et al., 2012), theta/alpha (Daume et al., 2017) and beta/gamma (Axmacher et al. 2008) frequency ranges have been observed to facilitate WM maintenance (for review see Fell and Axmacher, 2011). Furthermore, inhibiting the prefrontal cortex via transcranial magnetic stimulation has been shown to reduce WM encoding-related phase synchronization between the right frontal and central posterior electrode sites in the alpha frequency range (Zanto et al., 2011). Marshall et al. (2015) found that attention-dependent modulations of alpha power over the parietal cortical areas are related to fronto-parietal white-matter volume, suggesting that local parietal alpha activity relies on top-down influences from prefrontal cortex (PFC).

However, it remains elusive if alpha phase coherence between prefrontal cortex and the visual cortical areas supports WM maintenance of task-specific information. The alpha inhibition model posits that not only local alpha power (Klimesch, 2012; Jensen et al., 2012; 2014; Roux and Uhlhaas, 2014) but also long-range phase connectivity may support functional inhibition (Jensen and Mazaheri, 2010; Bonnefond et al., 2017). This is important as it leads to the counterintuitive prediction that local alpha power decreases rather than increases are accompanied by long-range alpha phase synchronization during functional disinhibition of task relevant networks. However, these effects have not been demonstrated yet.

To test these hypotheses, we recorded intracranial EEG activity in presurgical epilepsy patients performing a delayed-matching-to-sample task (as in Jokisch and Jensen, 2007) with three experimental conditions: maintenance of face identity (depending on the ventral visual stream, VVS), maintenance of face orientation (depending on the dorsal visual stream, DVS) and a control condition not requiring WM. We leveraged the high spatial and temporal resolution of the intracranial EEG recordings to answer three questions: 1) Does alpha activity support functional inhibition in the ventral and dorsal visual streams? We predicted reduced alpha activity during maintenance of face identity vs. control in the VVS and during maintenance of face orientation vs. control in DVS. 2) Is this effect associated with changes in the duration of duty cycles? We hypothesized longer duty cycles for identity vs. control in the VVS and for face orientation vs. control in the DVS. 3) Do alpha phase-based interactions support maintenance of relevant information? We predicted increased prefrontal-VVS alpha synchronization during maintenance of face identity vs. control and increased prefrontal-DVS alpha synchronization during maintenance of face orientation vs. control.

## Results

### Behavioral results

We analyzed behavioral accuracy (percentage of correct trials) for the three experimental conditions (see Figure 1A-C). A rank based nonparametric Kruskal-Wallis test (which was applied because the variance were non-homogenous; see Methods) with condition (orientation, identity, control) as factor revealed a significant main effect (χ^2^(2) = 11.69, p = 0.002; see Figure 1D). Post-hoc Wilcoxon signed-rank tests showed that accuracy in both the orientation (median ± standard deviation, 91.6% ±7.5%; Z = 2.57, p = 0.009) and identity condition (86.9% ±9.6%; Z = 2.58, p = 0.009) was lower than in the control condition (97.6% ± 8.1%), while accuracy did not differ between the identity and orientation conditions (Z = 1.49, p = 0.13). This suggests that the WM task was indeed more demanding in both the orientation and identity conditions as compared to the control condition, and that the identity and orientation conditions were similarly difficult.

**Figure 1:**
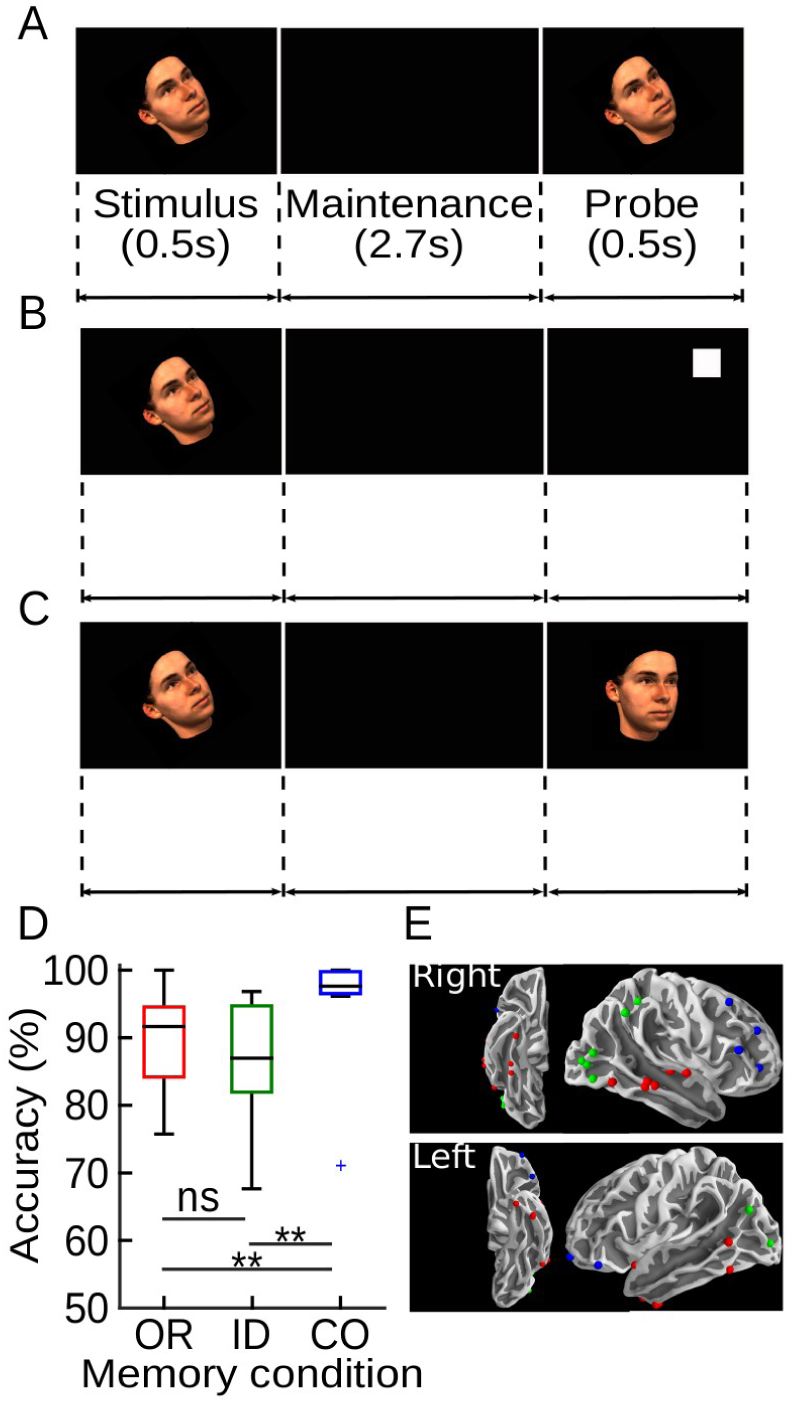
Experimental paradigm, behavioral results and electrode layout. ***(A-C)*** Trial structure with example of memory stimulus, maintenance interval and a matching probe during maintenance of identity ***(A),*** orientation ***(B),*** and right orientation probe in the control condition ***(C)***. Depending on condition, participants were instructed to encode and maintain the identity (ID) or orientation (OR) of the stimulus or to view it passively (control, CO). The same set of stimuli was used across three conditions – only instructions and the probe differed. ***(D)*** Accuracy scores as a function of experimental condition. The central line reflects the median. The edges present 25^*th*^ and 75^*th*^ percentile. Whiskers extend to the most extreme data points. ***(E)*** Inferior and lateral overviews of the selected electrodes in MNI space. Each dot represents an electrode from a single patient. Different colors indicate distinct regions of interests with green, red and blue corresponding to the dorsal and ventral visual stream and the prefrontal cortex, respectively. ** indicates p < 0.01.

### Double dissociation of alpha power reductions in feature-specific networks in ventral and dorsal visual stream

We analyzed intracranial EEG data from the ventral visual stream (VVS; N = 12 subjects) and dorsal visual stream (DVS; N = 7 subjects; Figure 1E) to investigate feature specific alpha power decreases. One electrode for each cortical region and for each patient was selected based on our localization procedure (see Methods). Subsequently, we performed label-shuffeled surrogate cluster analyses which controlls for multiple comparisons over frequency and time points (Maris and Oostenveld, 2007). In VVS, these analyses revealed significant reductions of alpha power during maintenance of face identity compared to either control (p = 0.03) or orientation (p = 0.02; Figure 2A-C). In contrast, in the DVS we found significant reductions of alpha power during maintenance of orientation compared to either control (p = 0.01) or identity (p = 0.01; Figure 2D-F). These results demonstrate a double dissociation of alpha power reductions during selective maintenance of task-relevant visual features in VVS and DVS. The observed effects were spectrally limited to the alpha frequency range (see Figure 2). This shows that alpha power decreases play a similar role in the ventral and dorsal visual streams; i.e. there is a relative decrease in alpha power in the stream that is engaged in the task and an increase in the stream that is not engaged.

**Figure 2:**
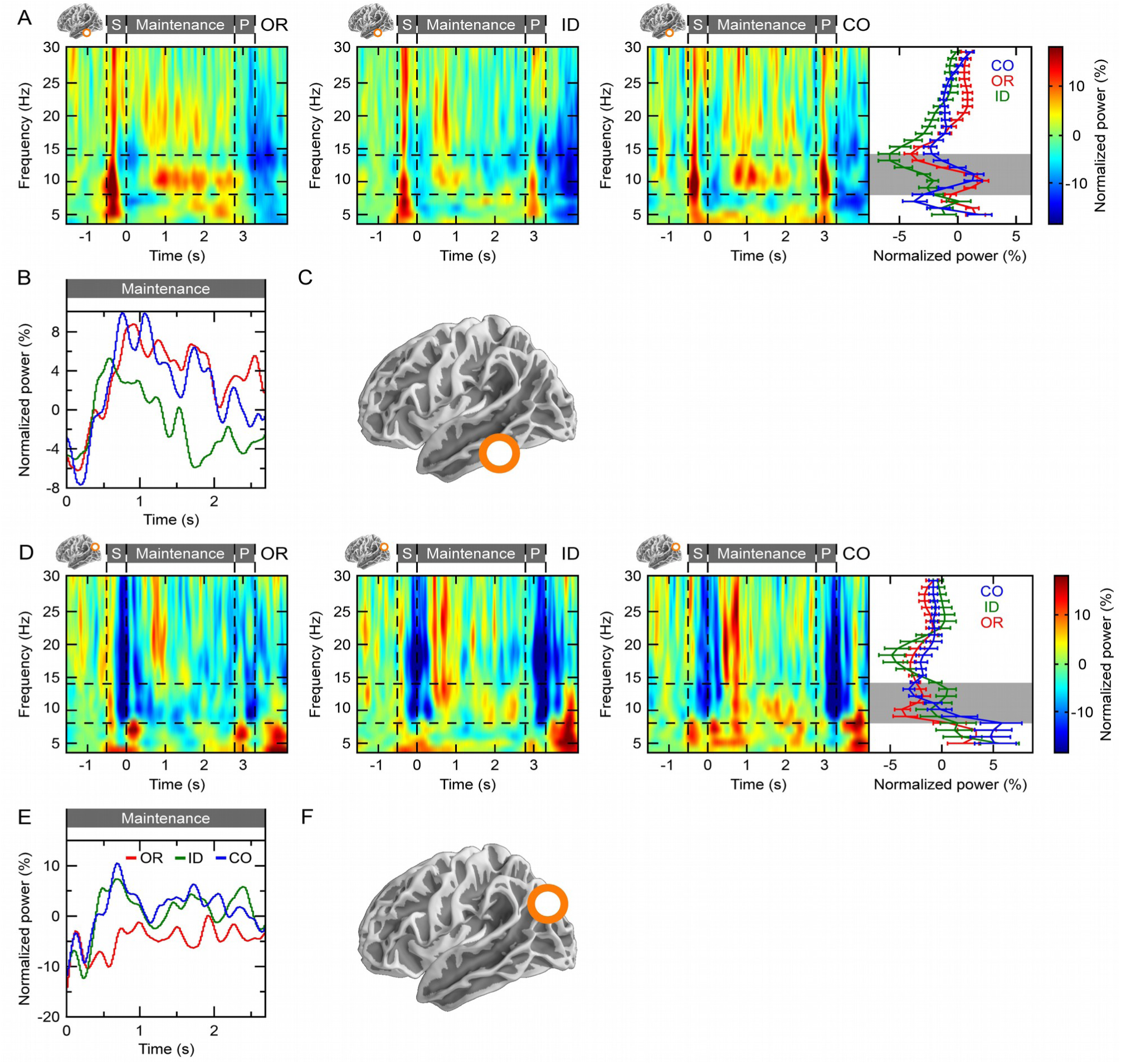
Feature specific alpha power reductions in ventral and dorsal visual stream. Low frequency (4-30Hz) activity patterns measured in ventral (A-C) and dorsal (D-F) visual stream (N = 12 and N = 7, respectively). We selected one electrode per region of interest in each subject, using a procedure that was independent of subsequently tested hypotheses (see Methods). (A, D) The time-frequency representations of power are plotted separately for the three experimental conditions (OR: Orientation, ID: identity and CO, control). Gray bars above figures indicate the trial structure consisting of stimulus presentation (S), maintenance, and probe presentation (P). Color indicates power normalized relative to the prestimulus baseline period, i.e. percentage change relative to average baseline power (B, E) Feature-specific alpha (8-14Hz) power reduction during the maintenance interval in ventral and dorsal visual stream, respectively. (C, F) Schematic illustration of the ventral and dorsal ROI, respectively.

### Alpha-gamma phase-amplitude coupling during WM maintenance

As a first step, we assessed overall (i.e. condition-independent) cross-frequency coupling (CFC) between the phase of alpha oscillations and the power of broad-band gamma activity (Roux and Uhlhaas, 2014). Broad-band high frequency activity (31-150 Hz) was selected because it has been suggested to reflect local neural activity (Mukamel et al., 2005; Ray et al., 2008). In each single patient, and in both VVS and DVS, alpha-gamma CFC in the empirical data was significantly higher than in trial-shuffled surrogates, with higher broadband power during the peaks than troughs of alpha oscillations (Figure 3A-B, C-D for the VVS and DVS, respectively). The distribution of surrogates was created by calculating CFC between alpha and broad-band gamma from randomly shuffled trial pairs. This procedure preserves the characteristics of the time-series and only eliminates a potential coupling between low and high-frequency time-series in individual trials (see Methods for more details). The overall magnitude of CFC did not differ across condition in the VVS (paired samples t-test: all t < 0.81; all p > 0.43) and in the DVS (all t < 1.28; all p > 0.44). The mean frequency of the phase-modulated gamma activity was at 76 ± 28 Hz (mean ± standard deviation) in VVS and at 92 ± 35 Hz in DVS with no difference across the two cortical areas (independent samples t-test: t(17) = 1.15; p = 0.26).

**Figure 3:**
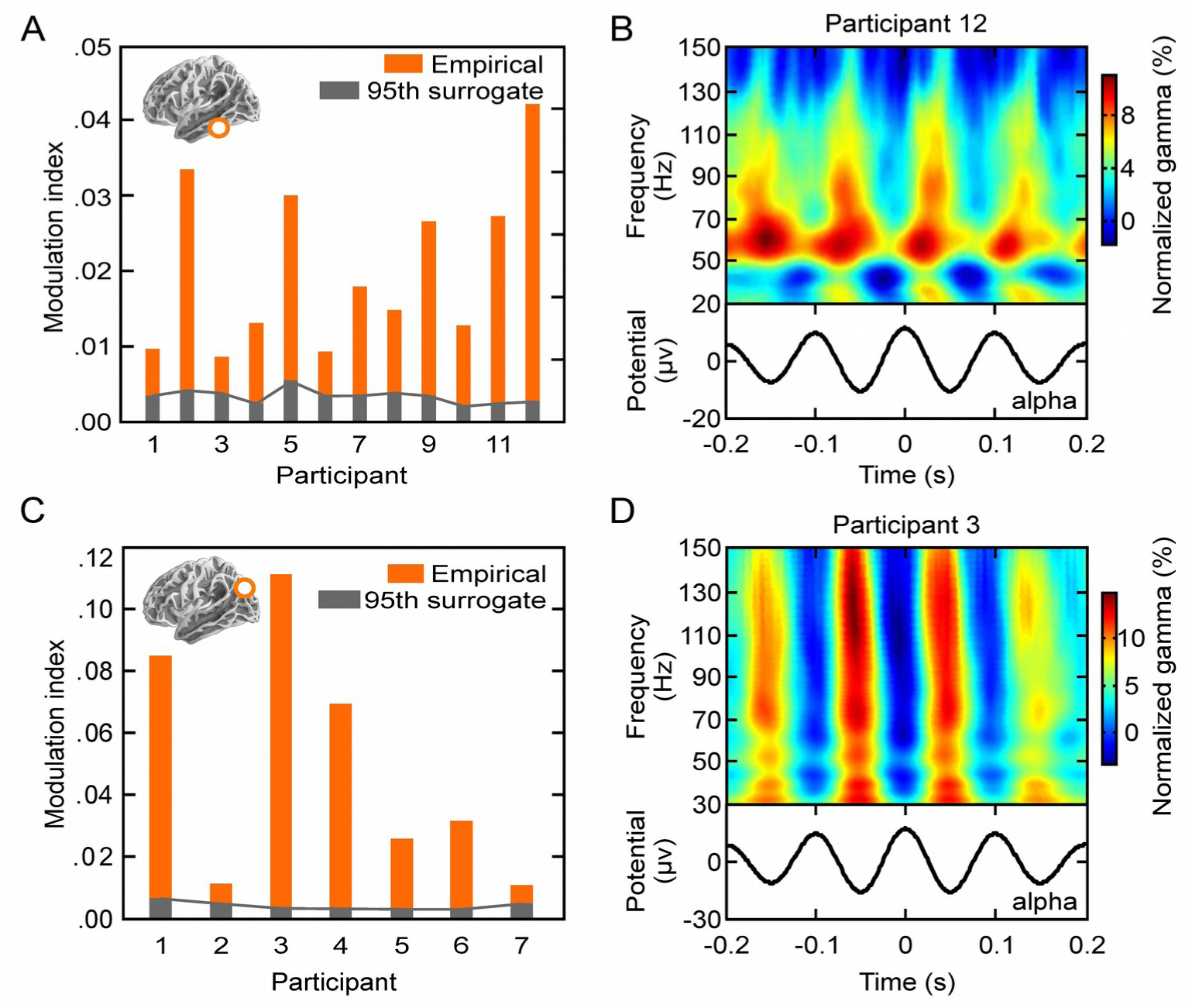
Alpha-gamma cross frequency coupling code. (A, C) In all subjects with electrodes in ventral (A) and dorsal visual stream (C), broadband high-frequency activity (31-150Hz) was significantly modulated during a specific phase (i.e. the peak) of alpha activity. Each subject showed increases of cross-frequency coupling (orange bars) above the 95^*th*^ percentile of a surrogate distribution (gray bars), corresponding to p < 0.05. Insets present schematic location of the ROI. (B, D) Individual examples of alpha-phase to gamma amplitude cross-frequency coupling in ventral (B) and dorsal (D) visual stream. Color represents normalized broadband gamma (31-150Hz) created by subtracting and dividing gamma amplitude values by the mean amplitude across the time interval to facilitate visualization across frequency bands.

### WM maintenance of relevant features is associated with longer duty cycles

Next, we tested if reductions of alpha power are related to longer “duty cycles” in feature-specific brain areas (Jensen et al., 2012, 2014; Leszczynski et al., 2015). Duty cycles are reflected by specific alpha phases during which broad-band gamma activity is relatively enhanced. The alpha phase at which gamma activity is maximal can be specified by calculating the modulation phase (Canolty et al. 2006). In case of longer duty cycles the maximum of gamma activity may be distributed across a wider range of alpha phases. Thus, we hypothesized that the circular variance (Fisher, 1996) of modulation phases across trials should be increased in a given area during WM maintenance of relevant features compared to control. To this end, we quantified the circular variance of the alpha-gamma CFC modulation phases for the three experimental conditions in both VVS and DVS. Indeed, modulation phases were more disperse in the respective task-relevant condition in VVS (t-test; identity vs. control condition: t(11) = 2.29, p = 0.04) and DVS (orientation vs. control: t(6) = 2.57, p = 0.04; see Figure 4). No other differences were observed (all t < 1.72, all p > 0.11). To exclude the possibility that these effects result spuriously from differences in signal-to-noise ratio due to condition differences in alpha power, we compared the inter-trial phase coherence across experimental conditions. We reasoned that any condition specific differences in the reliability of phase extraction would result in differences in coherence. However, the magnitude of inter-trial phase coherence did not differ between experimental conditions (all t < 0.44; all p > 0.66). This excludes the possibility that the condition differences in duty cycles are only due to differences in signal-to-noise ratio affecting the reliability of phase extraction. Importantly, trial numbers did not differ between conditions (CO vs. ID: t(11) = 1.04, p = 0.32; CO vs. OR: t(6) = 1.25, p = 0.26) which makes it also unlikely that unequal number of trials explains the observed differences in the circular variance. Nevertheless, we performed a further control analysis in which we confirmed that the results hold with trial numbers exactly matched. To this end, we randomly selected trials from the condition with the larger number of trials in each participant such that it matched the number of trials in the condition with fewer trials. This procedure was repeated 100 times and the average modulation variance across these 100 samples (with exactly matched trial numbers) was used for comparisons. Again, we observed that modulation phases were more disperse in the task-relevant condition in both VVS and DVS (VVS, identity vs. control condition: t(11) = 2.69, p = 0.02; DVS, orientation vs. control: t(6) = 2.46, p = 0.04). No other differences were observed (all t < 1.53, all p > 0.15).

**Figure 4:**
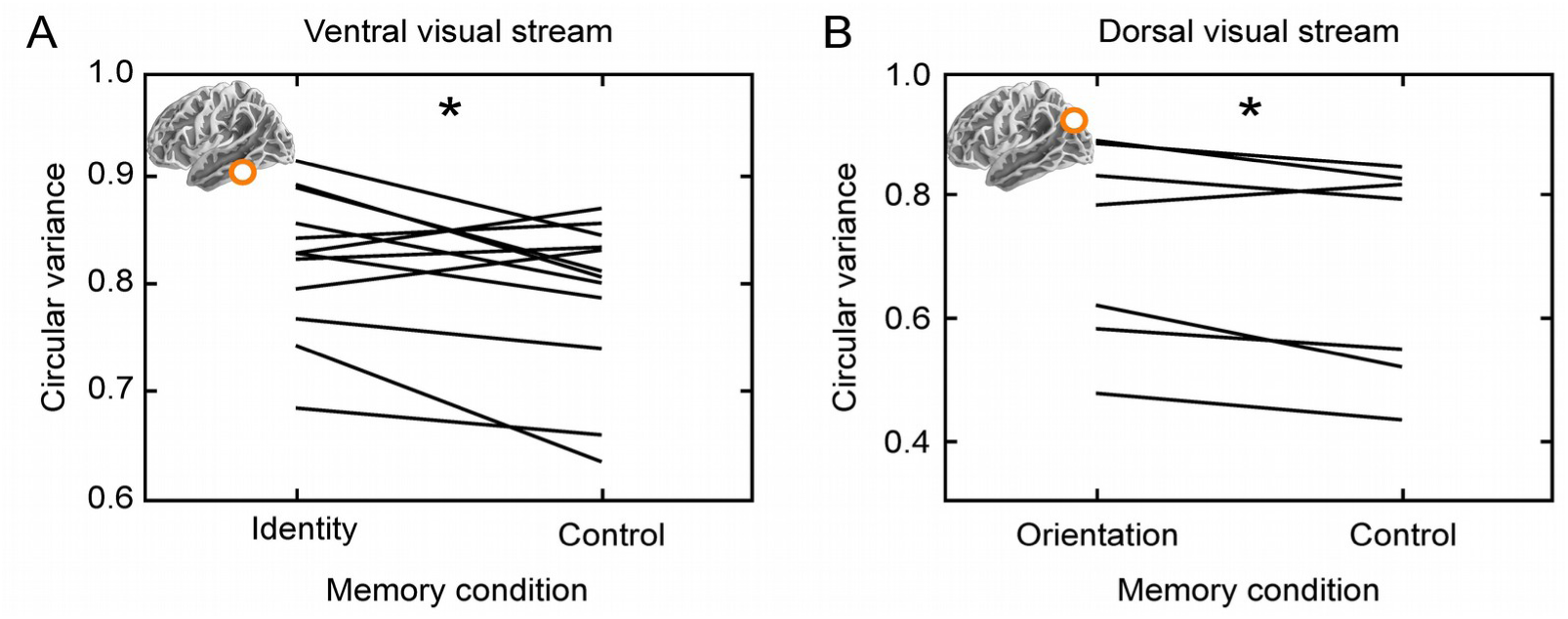
Broader duty cycles during maintenance of task-relevant features. Circular variance (calculated across trials) of modulation phases (high-frequency power across alpha phase) in ventral ***(A)*** and dorsal ***(B)*** visual stream. Each line represents a single subject (N = 12 and N = 7 for the ventral and dorsal visual stream, respectively). Insets represent the ROIs. * indicates p < 0.05.

In sum, our data show that gamma activity occurs during more disperse alpha phases when relevant features are being maintained. This is in line with the hypothesis that decreasing alpha power enhances information processing by increasing the length of excitable duty cycles.

### Phase synchronization between prefrontal cortex and feature specific visual networks

In some patients electrodes were implanted simultaneously in VVS and PFC (N = 5) or in DVS and PFC (N = 2). We used these simultaneous recordings to investigate whether the ventral and dorsal visual stream differentially interacted with prefrontal cortex during maintenance of relevant features. To this end, we performed a functional connectivity analysis based on alpha phase synchronization. All five patients with electrodes in both VVS and PFC showed increased phase synchronization during maintenance of relevant (i.e., face identity) and irrelevant (i.e. face orientation) features compared with control (see Figure 5A-B). Because of the relatively small groups of patients, we did not conduct group statistics. Both patients with electrodes in DVS and PFC showed the opposite pattern, i.e. increased phase synchronization during maintenance of orientation vs. control (see Figure 5C-D). Notably, these condition differences are in opposite direction to the changes in alpha power, so that higher phase synchronization cannot be explained by increased alpha power. No other condition differences were consistent. This supports the hypothesis that alpha phase synchronization between PFC and task-relevant visual areas contributes to maintenance of information in WM. These results although informative should be treated with caution until reproduced with larger sample sizes.

**Figure 5:**
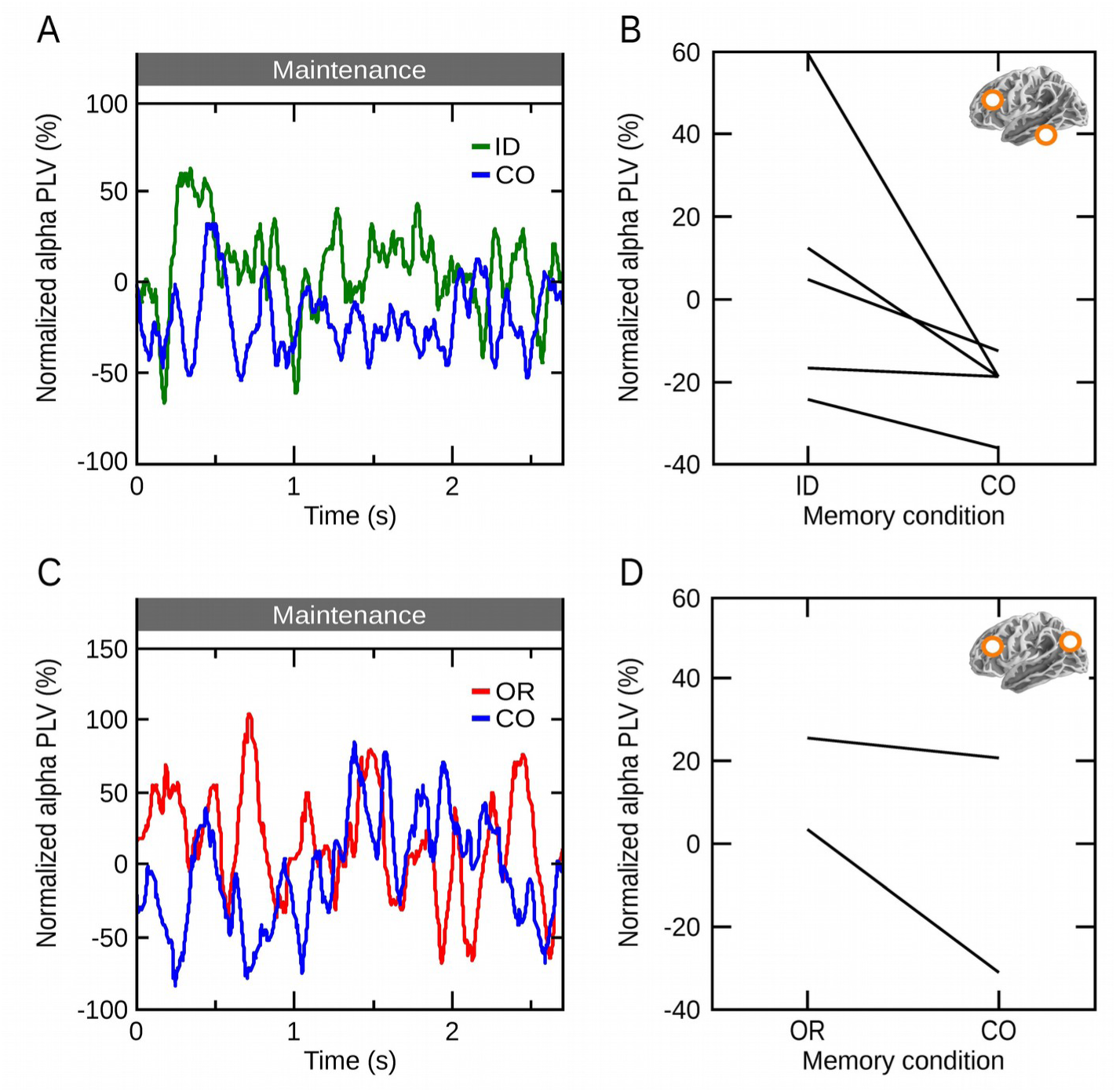
Increased alpha phase synchronization with prefrontal cortex during maintenance of task-relevant features. Alpha phase synchronization during maintenance between prefrontal cortex and ventral (A, B) as well as dorsal (C, D) visual stream averaged across patients (N = 5, N = 2, for ventral visual stream, dorsal visual stream and prefrontal cortex, respectively) normalized relative to the prestimulus baseline interval. (B, D) Alpha phase synchronization averaged across the entire maintenance interval for each participant. Lines indicate single subjects. Insets represent the ROIs. Due to the low number of subjects with these electrode combinations we did not conduct group statistics.

## Discussion

### Alpha activity shows a double dissociation between feature specific networks in ventral and dorsal visual stream

In the current study, we leveraged the high temporal and spatial resolution of intracranial EEG recordings to demonstrate that maintenance of either the identity or spatial orientation of the same face stimuli is achieved by selective decreases of alpha power in the ventral or dorsal visual stream, respectively. By contrast, alpha power remained strong in the stream that was not engaged by the tasks. This pattern of results supports the notion that alpha oscillations serve a role in the active functional inhibition of areas not required for a given task (Jokisch and Jensen, 2007; Foxe and Snyder, 2011; Klimesch, 2012; Bonnefond and Jensen, 2012; Jensen et al., 2012; Jensen et al., 2014; Anderson and Hanslmayr, 2015). While previous studies tested this idea mostly in primary sensory areas (Jokisch and Jensen, 2007; Stokes et al., 2009; Haegens et al., 2011; Spaak et al., 2012; Payne et al., 2013), the current results extend our knowledge by demonstrating that alpha power also supports functional inhibition of higher order visual areas. These results are particularly noteworthy because previous studies have suggested different physiological mechanisms and functions of alpha activity in VVS as compared to early sensory cortices (Bollimunta et al., 2008; Mo et al., 2011). The current results suggest that the functional role of alpha oscillations generalizes across extended visual areas including the ventral and dorsal visual stream.

### The alpha-gamma code in the neocortex

In addition to the alpha amplitude changes, we showed task dependent modulations of the duty cycle (i.e. the distribution of alpha phases associated with a high power of broadband high-frequency activity). In particular, we found that duty cycles were more broadly distributed across trials when relevant features were being maintened, both in ventral and dorsal visual stream (see Figures 4 and 6). This supports a model in which duty cycles are defined as intervals during which neural activity is released from pulses of alpha inhibition (Haegens et al., 2011, Spaak et al., 2012; Samaha and Postle, 2015; for review see Jensen et al., 2014; VanRullen, 2016). Furthermore, the current results suggest a relation between alpha amplitude and the duration of the duty cycle, with lower alpha amplitudes corresponding to longer duty cycles (Jensen et al., 2014; see Figure 6).

**Figure 6:**
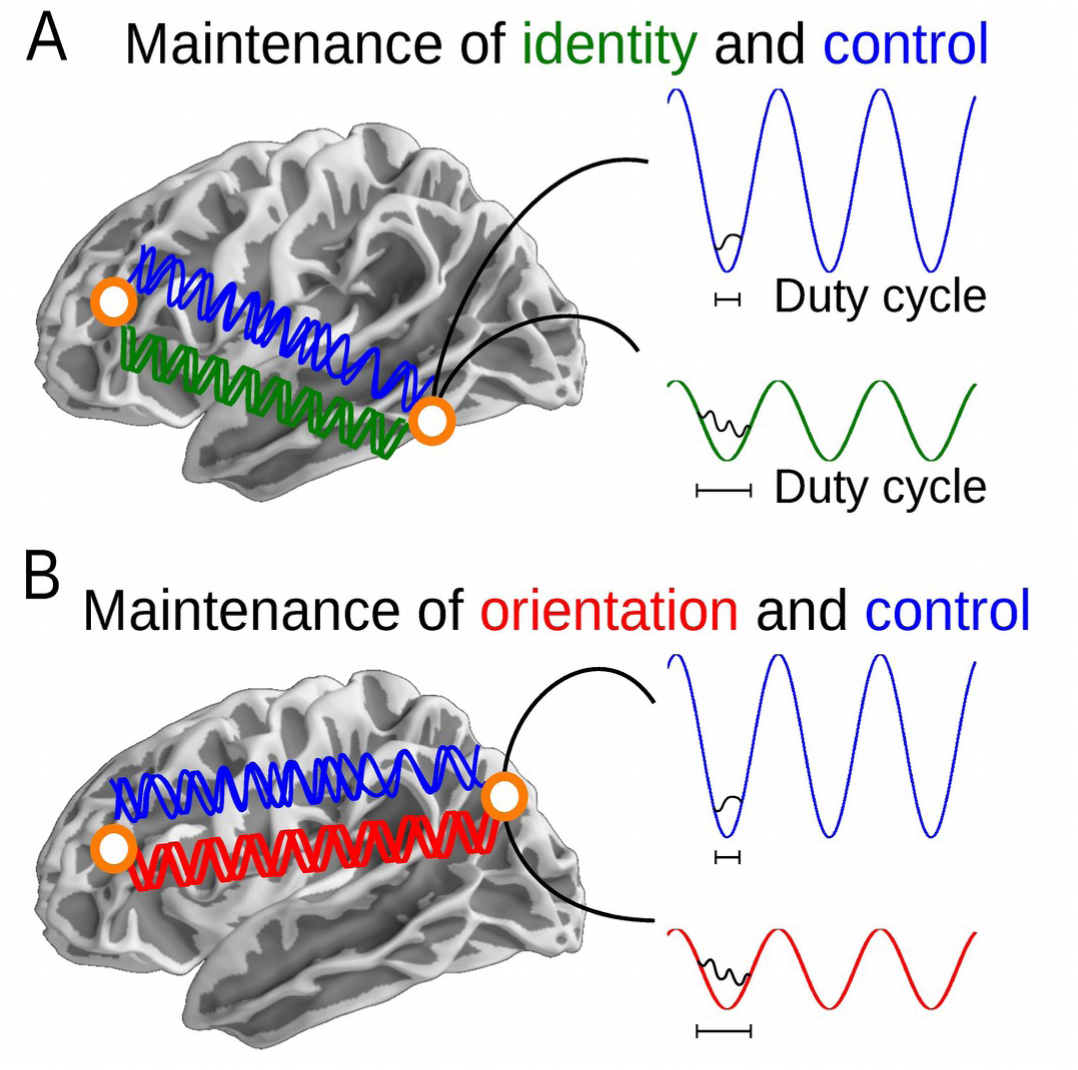
Model of functional inhibition due to local and long-range coordination of alpha oscillations. ***(A,B)*** Alpha power changes and alpha phase synchronization in the ventral visual stream and prefrontal cortex ***(A)*** and dorsal visual stream and prefrontal cortex ***(B)*** networks. Plots are illustrative of significant results. Lower alpha power during maintenance of task-relevant features (i.e. face identity and orientation in ventral and dorsal visual stream, respectively) is accompanied by more variable (i.e. extended duty cycles during which high-frequency gamma activity is increased). Local alpha power reductions also coincide with stronger phase-based network connectivity between the respective visual areas and the prefrontal cortex.

In general, we observed significant cross-frequency coupling between alpha phase and broadband high-frequency power in both VVS and DVS. The magnitude of this coupling was not modulated by the task. These findings suggest that the duty cycle of high-frequency activity with regard to alpha phase rather than the coupling strength itself reflects task demands. This is consistent with the cross-frequency coupling model of WM, which relates the representation of individual items in WM to high-frequency activity distributed across low-frequency oscillations (Lisman and Idiart, 1995; Lisman and Jensen, 2013; Roux and Uhlhaas, 2014). The current results together with previous findings from the hippocampus (Leszczynski et al., 2015; Heusser et al., 2016) and neocortex (Vaz et al., 2017) during working and episodic memory suggest that the distribution rather than the magnitude of high-frequency activity supports successful maintenance of relevant information.

It has been convincingly shown that artificial cross-frequency coupling modulations might be observed due to changes in power (Aru et al., 2015) or the shape of the signal (Cole and Voytek, 2017; but see Jensen et al., 2016). Here, we observed no difference in the CFC magnitude despite modulations of the underlying alpha power. This suggests that even though alpha power was reduced, it still exercises a phasic modulation of the high broadband activity. This also reduced concerns on spurious CFC coupling.

### The long-range network connectivity

Because simultaneous multi-cortical implantations tend to be rare, there was a limited number of patients with recordings in both prefrontal cortex/VVS and prefrontal cortex/DVS (N = 5 and N = 2, respectively). Therefore, the current long-range network connectivity results should be interpreted with caution until corroborated by further evidence. We used multi-cortical recordings in the VVS, DVS and PFC to study the neural mechanism supporting interactions across these cortical nodes. This is important because prefrontal cortex and sensory cortices have been put forward as core regions of the neural architecture for WM (Curtis and D’Esposito, 2003; Sreenivasan et al., 2014; Eriksson et al., 2015). We observed that the cortical region involved in representing a particular visual feature showes increased alpha phase connectivity with the prefrontal cortex (5 subjects out of 5 for PFC-VVS and 2 subjeects out of 2 for PFC-DVS). Previous electrophysiological work has demonstrated that increased synchrony in theta (Liebe et al., 2012), theta/alpha (Daume et al., 2017), alpha (Sauseng et al., 2005a,b, 2011; Popov et al. 2017) and beta/gamma (Axmacher et al. 2008) supports WM representation. Furthermore, alpha power over parietal sites relates to fronto-parietal white matter volume (Marshall et al., 2015). Here, we extend these previous results by showing increased alpha phase connectivity with the cortical areas relevant for maintenance of a feature (i.e. ventral and dorsal visual stream for identity and orientation, respectively). Importantly, these cortical areas show decreased alpha power during the very same conditions. This is relevant because it supports a model in which increased local alpha amplitude facilitates information propagation by blocking redundant pathways (Bonnefond et al. 2017). Second, it suggests that alpha oscillations remain functionally relevant even when local power is low (i.e. the decreased alpha power enables formation of cortical ensembles). This indicates that alpha shapes both local and long-range functional architecture.

### WM is implemented in a distributed network

WM is a key process underlying various cognitive functions. It has been suggested to rely on a distributed network involving several cortical areas (Eriksson et al., 2015). Previous fMRI studies identified distributed cortical regions supporting WM maintenance (for review see Christophel et al., 2017) and pointed towards ventral and dorsal visual stream as relevant for maintenance of face identity and orientation. Here we extend these results showing that local-and long-range alpha activity is the mechanism supporting flexible maintenance of distinct information pieces in WM. Together, our findings highlight the role of alpha dependent coordination between ventral, dorsal visual stream and the prefrontal cortex which operate together for efficient and flexible WM maintenance.

## Materials and Methods

### Participants

EEG was recorded from 12 pharmacoresistant epilepsy patients (mean age of 32.2 years, range 18 to 55 years, six female) implanted with subdural grid and stripe electrodes for presurgical diagnostic purposes. All patients had well-defined ictal onset zones. Eight of them had a seizure onset zone within the left hemisphere, the other three in the right hippocampus and one had a bilateral hippocampal onset zone. Only electrodes that were located outside of the focus of epilepsy were considered for analysis. The patients had electrodes located in the ventral visual stream (VVS, N = 12) and dorsal visual stream (DVS, N = 7) as well as in the prefrontal cortex (PFC, N = 5). Five patients had simultaneous recordings in VVS and PFC, and two had electrodes in both DVS and PFC. Recordings were performed at the Department of Epileptology, University of Bonn, Bonn, Germany. The study was approved by the local medical ethics committee, and all patients gave written informed consent. Minimal requirement was the ability to successfully complete the experiment and availability of the pre-and post-implantation MRI scans to localize electrodes.

### Task

Participants performed a delayed matching to sample WM task which was designed to engage either the ventral or dorsal visual stream (see also Jokisch and Jensen, 2007). Depending on the experimental condition, participants maintained the identity of a face, its spatial orientation, or passively viewed the face (“Identity”, “Orientation” and “Control” conditions, see Figure 1A-C). We reasoned that maintenance of the same briefly presented faces will involve either the ventral (Courtney et al., 1997; Druzgal and D’Esposito, 2003; Postle et al., 2003; Ranganath et al., 2004a,b) or dorsal (Awh et al., 1999; Postle and D’Esposito, 1999; Postle et al., 2004; Postle, 2006) visual stream depending on the feature the participants were instructed to maintain (i.e. face identity or orientation, respectively). We furthermore expected increased network connectivity with the PFC during maintenance of the relevant feature because the PFC has been suggested to exert top-down control on the sensory cortical areas (Curtis and D’Esposito, 2003; Sreenivasan et al., 2014).

Each trial started with the presentation of an image of a face for 0.5s, followed by a maintenance interval lasting 2.7s. Next, participants were presented with a second stimulus (0.5s) which probed their WM for specific features of the face (see Figure 1A-C). The probe stimulus was either a face (identity and control conditions) or a box (orientation condition) and could either match the maintained feature (face identity or its orientation depending on the condition) or not with equal probability. Faces were presented at 12 different angles of rotation (-60°, −30°, 30°, 60° relative to the vertical axis and −30°, 0, 30° relative to horizontal axis, see Jokisch and Jensen, 2007). Participants performed a two alternative forced choice task, indicating for each trial whether the probe did or did not match the memorized feature. After the response, participants were provided with feedback regarding accuracy in a given trial. During the control condition participants were instructed to passively view the image of the first face and to indicate if the image of the second face was rotated to the left or to the right. The visual input and the response requirements were similar across both memory conditions and control, so that perceptual features could not explain differences in task-related activity. However, the control condition did not involve any memory load because participants’ reactions only depended on the probe stimuli. The order of conditions was pseudo-randomized across patients. Participants were instructed to respond as accurately as possible.

### Electrophysiological recordings

The data was recorded using multicontact strip and grid electrodes. These electrodes were made of stainless steel and had contacts with a diameter of 4mm and an inter-contact center-to-center spacing of 10mm. All data was referenced to the linked mastoids, on-line band pass filtered between 0.01Hz (6dB/octave) and 300Hz (6dB/octave) and then sampled at 1kHz.

### Data processing

All data were processed offline using MATLAB (MathWorks) as well as the Circular Statistics Toolbox (Berens, 2009). Prior to processing, all data from electrodes clinically identified to be within the ictal onset zone were removed. All data pre-processing was performed at the single subject and single electrode level. For each subject the data from all non-ictal electrodes were notch-filtered to remove 50 Hz line noise and its harmonics. The data was subsequently visually inspected for any remaining artifacts (predominantly arising from the epileptiform activity). All samples contaminated by artifacts were marked and removed from further analysis.

### Regions of interest

We selected one electrode per patient in each region of interest (ROI; VVS, DVS and PFC) based on a combination of anatomical and functional characteristics (for a similar approach, see for example Axmacher et al., 2010a; Leszczynski et al., 2015; Voytek et al., 2015; Foster et al., 2015; Cogan et al., 2017). All selection criteria were orthogonal to the subsequent analyses. Pre-and post-operative MR images were used to classify electrodes into the three ROIs. The electrode localized in the lateral temporal lobe which showed the largest intracranial face selective N200 ERP component was classified as VVS site (Allison et al., 1999, McCarthy et al., 1999). This criterion is orthogonal to all subsequent analyses because it is based on the signal during face presentation across all three conditions. The most posterior electrode in the region of the superior occipital or parietal lobe was selected as DVS electrode. This was based on the activation pattern observed by Jokisch and Jensen (2007; i.e. alpha power changes were most prominent over occipital sites extending to the parieto-occipital sulcus) using a very similar paradigm with the same stimuli. Finally, one electrode anterior to the precentral sulcus with the largest sustained DC shift during retention was selected as PFC channel (Curtis and D’Esposito, 2003; Sreenivasan et al., 2014).

### Behavioral analysis

To estimate behavioral performance we calculated accuracy scores (percentage of correct trials) separately for each condition and participant and compared them using a Kruskal-Wallis test and post-hoc non-parametric Wilcoxon signed rank test (Hollander and Wolfe, 1973). This approach rather than analysis of variance was used because the behavioral data violated assumption of homoscedacity. Kolmogorov-Smirnov and Levene’s tests were used to test assumptions of normality and homoscedacity.

### Time-frequency analysis

Continuous EEG data was segmented into 8s long epochs with a 2s pre-stimulus period and a 6s post-stimulus interval relative to the presentation onset of the memory stimulus in each trial (see Figure 1A). Such relatively long segments were used to minimize edge effects. Only artifact free segments during correct trials were considered for analyses. The epoched data was convolved with seven-cycle Morlet wavelets from 3Hz to 30Hz in steps of 1Hz using functions from Fieldtrip Matlab toolbox (Oostenveld et al., 2011). The power values were normalized with respect to the prestimulus time windows from −0.7s to −0.2s. Only retention related activity during the time period after the offset of the cue and across the whole 2.7s maintenance period was further analyzed. We used a non-parametric label-shuffled surrogate statistics with cluster correction for multiple comparisons over time and frequency (Maris and Oostenveld, 2007) to test our a priori hypothesis of decreased alpha power during maintenance of the relevant features both in the ventral and dorsal visual stream. In this initial step we targeted the whole alpha spectrum (8-14Hz). For further analysis we focused on the peak alpha frequency where the permutation-based cluster controlled t-statistics showed the most prominent difference in the grand average between maintenance of relevant and control features (i.e., 12Hz in the VVS and 10Hz in the DVS; these frequencies were not significantly different between streams).

### Analysis of cross-frequency coupling

Epoched data was convolved with seven-cycle Morlet wavelets from 3Hz to 150Hz in steps of 1Hz. Cross-frequency coupling was analyzed using a previously described procedure (Axmacher et al., 2010b; Leszczynski et al., 2015) based on Pearson’s correlations between the signal in the alpha frequency band (obtained via wavelet convolution) and the power time series of the high-frequency signal (30-150Hz; for a similar method, see the “envelope-to-signal correlation” described by Penny et al., 2008; Tort et al., 2010). To account for possible inter-individual differences between the phases of the low-frequency signal at which the amplitude of the high-frequency signal was maximal, the low-frequency signal was shifted by the subject-specific “average modulation phase” at which the high-frequency amplitude was maximal for each participant. In details, we first extracted low- and high-frequency signals with seven-cycle Morlet wavelets, resulting in complex-valued time-series for low-frequency (activity-low) and high-frequency (activity-high) signals. The phase of low-frequency signals (phase-low) and the amplitude of high-frequency signals (amplitude-high) were used to construct a composite signal z = amplitude-high * e^i*phase-low^ (Canolty et al., 2006). Subsequently, the low-frequency signal was shifted by the average modulation phase. Finally, Pearson’s correlation between the real part of the phase-shifted low-frequency signal and the amplitude of the high-frequency signal was calculated.

To estimate if the observed alpha-gamma modulation during the maintenance interval was higher than chance, we compared this empirical CFC to a reference distribution of surrogate data obtained by randomly re-assigning trials for amplitude and trials for phase. This was done separately in each participant. All consecutive analysis steps were exactly the same as for the calculation of the empirical data. The procedure was repeated 100 times for each subject, resulting in a reference distribution under the null hypothesis that there is no phase-amplitude coupling. Importantly, this procedure for calculating surrogates preserves the analytic amplitude and phase time series and only randomizes the relative trial structure between the two variables. This non-parametric method allows for testing whether the observed CFC is larger than the reference distribution. To this end, we sorted the surrogate CFC values and accepted the empirical CFC as significant if it exceeded the value of the 95^th^ percentile which corresponds to a threshold of p < 0.05 (for similar methods see Lachaux et al., 1999; Voytek et al., 2013).

### Analysis of modulation variance

To quantify the variance of the position of the phase-modulated high-frequency signal with respect to alpha phase, the complex-valued composite signal (z = amplitude-high * e^i*phase-low^) was first averaged across the maintenance interval in each trial. Then, the single-trial modulation phase value was extracted (i.e., the alpha phase at which broad-band gamma power was maximal in a given trial). The circular variance was calculated across the distribution of trial-wise modulation phases separately for each condition and each participant (Fisher, 1996; Berens, 2009). This analysis reflects the consistency of the alpha phase at which the high frequency amplitude is increased. The higher the variance, the broader the distribution of gamma power across the alpha “duty” cycle. Variance was compared to control condition using t-test.

### Analysis of phase synchronization

We calculated phase synchronization between PFC and VVS as well as between PFC and DVS during the whole maintenance interval. To this end, we calculated the circular variance of phase differences (Lachaux et al., 1999) separately for signals between prefrontal - ventral visual stream (N = 5) and prefrontal – dorsal visual stream (N = 2) sites. The method quantifies the stability of phase differences between two time series across trials. Phase synchronization was compared between WM conditions. Combined anterior and posterior implantations tend to be rare (N = 5 between PFC and VVS; N = 2 between PFC and DVS), so we refrained from group statistics and instead assessed whether the effects were consistent across all patients.

## Author Contributions

ML, JF, OJ and NA designed the study. ML analyzed the data. ML, JF, OJ and NA wrote the manuscript.

## Acknowledgments

The authors would like to thank Anne Do Lam for support during data collection, and Amirhossein Jahanbekam for helpful discussions. ML and NA were supported by an Emmy Noether grant by the DFG (AX82/2). NA received additional funding through SFB 874 and DFG project AX 82/3 and together with JF via SFB 1089.

